# Targeting pentose phosphate pathway for SARS-CoV-2 therapy

**DOI:** 10.1101/2020.08.19.257022

**Authors:** Denisa Bojkova, Rui Costa, Marco Bechtel, Sandra Ciesek, Martin Michaelis, Jindrich Cinatl

## Abstract

It becomes more and more obvious that deregulation of host metabolism play an important role in SARS-CoV-2 pathogenesis with implication for increased risk of severe course of COVID-19. Furthermore, it is expected that COVID-19 patients recovered from severe disease may experience long-term metabolic disorders. Thereby understanding the consequences of SARS-CoV-2 infection on host metabolism can facilitate efforts for effective treatment option. We have previously shown that SARS-CoV-2-infected cells undergo a shift towards glycolysis and that 2-deoxy-D-glucose (2DG) inhibits SARS-CoV-2 replication. Here, we show that also pentose phosphate pathway (PPP) is remarkably deregulated. Since PPP supplies ribonucleotides for SARS-CoV-2 replication, this could represent an attractive target for an intervention. On that account, we employed the transketolase inhibitor benfooxythiamine and showed dose-dependent inhibition of SARS-CoV-2 in non-toxic concentrations. Importantly, the antiviral efficacy of benfooxythiamine was further increased in combination with 2DG.

---

The rapid spread of SARS-CoV-2 resulted in millions of infected people worldwide. Though the majority of patients exhibit mild to moderate symptoms, around 20% of patients develop severe disease. Effective antiviral drugs against SARS-CoV-2 are currently lacking.

Emerging evidence suggests that metabolic disorders are associated with an increased risk of severe COVID-19. Increased amounts of glucose in the sera of COVID-19 patients were correlated with poor prognosis in individuals both with and without pre-existing diabetes (Shen et al., 2020). At the cellular level, SARS-CoV-2 induces a shift of the host cell metabolism towards glycolysis in infected cells, and the glycolysis inhibitor 2-deoxy-D-glucose (2DG), which targets hexokinase (the rate-limiting enzyme in glycolysis), interferes with SARS-CoV-2 infection in colon adenocarcinoma (Caco-2) cells and monocytes (Bojkova et al., 2020; Codo et al., 2020). Hence, agents that antagonize virus-induced changes in the host cell metabolism, especially glycolysis, are candidates for COVID-19 therapy.

Using a proteomics dataset derived from SARS-CoV-2-infected Caco-2 cells (Bojkova et al., 2020), we identified further metabolic changes associated with SARS-CoV-2 infection. Cellular levels of key drivers of glycolysis including hexokinase 1 and 2 (HK1 and HK2) were increased in SARS-CoV-2-infected cells, while negative regulators such as fructose-1,6-bisphosphatase 1 and 2 (FBP1 and FBP2) were reduced (Figure S1A). Changes in glycolysis can also affect other metabolic pathways. Metabolic intermediates generated by the glycolytic enzymes supply both the oxidative and the non-oxidative branch of the pentose phosphate pathway (PPP) with intermediates (Stincone et al., 2015). Notably, transketolase (TKT) and transaldolase 1 (TALDO1), two constituents of the non-oxidative PPP branch (Stincone et al., 2015), displayed increased levels in SARS-CoV-2-infected cells (Figure S1A), suggesting that the non-oxidative PPP branch is also regulated in response to SARS-CoV-2 infection.

The non-oxidative PPP branch converts glycolytic intermediates into ribose-5-phosphate, which is required for the synthesis of nucleic acids, and sugar phosphate precursors, which are necessary for the synthesis of amino acids (Stincone et al., 2015). TKT, a key enzyme of the non-oxidative PPP, becomes activated by the binding of one molecule of thiamine diphosphate and of a bivalent cation (Zhao and Zhong, 2009). Recently, it has been shown that TKT activity may be strongly increased by the formation of a heterodimer with transketolase like 1 (TKTL1) (Li et al., 2019).

Irreversible inhibitors of TKT such as oxythiamine (an inactive analogue of thiamine) have been developed (Tylicki et al., 2018). To investigate whether TKT inhibitors affect SARS-CoV-2 replication, we used benfooxythiamine (BOT), a prodrug of the TKT inhibitor oxythiamine, currently evaluated in pre-clinical experiments (Coy, 2017). Indeed, non-toxic BOT concentrations inhibited the replication of two different SARS-CoV-2 isolates (FFM1 and FFM7) in a concentration-dependent manner, as indicated by immunostaining for the SARS-CoV-2 Spike (S) protein and the detection of genomic RNA by qPCR (Figure S1B–S1D).

The antiviral effects of BOT were further increased by the glycolysis inhibitor 2DG (Figure S1D-SLF), indicating that the combined inhibition of ribose-5-phosphate production via the non-oxidative PPP branch and glycolysis is more effective in suppressing SARS-CoV-2 replication than the inhibition of either metabolic pathway alone. Mechanistically, 2DG inhibits the glycolysis enzymes hexokinase and phosphoglucose isomerase (PGI), both required for synthesis of fructose-6-phosphate, which serve as glycolytic intermediate for generating ribose-5-phosphate in non-oxidative PPP branch (Stincone et al., 2015).

Furthermore, ribose-5-phosphate can be also generated in the oxidative PPP branch through the metabolic conversion of glucose-6-phosphate (Stincone et al., 2015). Since 2DG decreases cellular levels of glucose-6-phosphate, it may additionally lower ribose-5-phosphate levels by reducing its availability for metabolizing in the oxidative branch of PPP. Importantly, the oxidative PPP branch seems also be activated in SARS-CoV-2-infected cells as indicated by increased 6-phoshogluconate dehydrogenase (PGD) levels (Figure S1A). PGD catalyzes the conversion of 6-phosphogluconate into ribulose-5-phosphate, a precursor of ribose-5-phosphate, in the oxidative PPP branch (Stincone et al., 2015).

Hence, 2DG could increase the effects of BOT by further reducing ribobose-5-phosphate synthesis through interrupting the supply of glycolytic intermediates into both oxidative and non-oxidative branch of PPP.

In conclusion, SARS-CoV-2 infection is associated with changes in the regulation of the PPP, which has potential implications for antiviral therapy. The TKT inhibitor BOT inhibited SARS-CoV-2 replication, indicating a role of the non-oxidative PPP branch in the virus replication cycle. The addition of the glycolysis inhibitor 2DG further enhanced the antiviral activity of BOT, which may result from the interruption of supply of glycolytic intermediates into PPP. The findings also demonstrate that the simultaneous inhibition of different metabolic pathways has potential as an antiviral treatment strategy for SARS-CoV-2-infected individuals (Figure S1G). Notably, uncontrolled glucose and pentose phosphate levels correlate with worse prognosis and higher mortality in COVID-19 patients (Deng et al., 2020; Thomas et al., 2020; Zhu et al., 2020), and reports show that a number of COVID-19 survivors suffer from metabolic disorders including abnormal glucose metabolism (Ayres, 2020). Therefore, targeting the host cell metabolism may prevent the metabolic derailment of immune cells in addition to the inhibition of SARS-CoV-2 replication and thus contribute to recovery in severe COVID-19 disease (Ayres, 2020).

---

**Figure S1.**
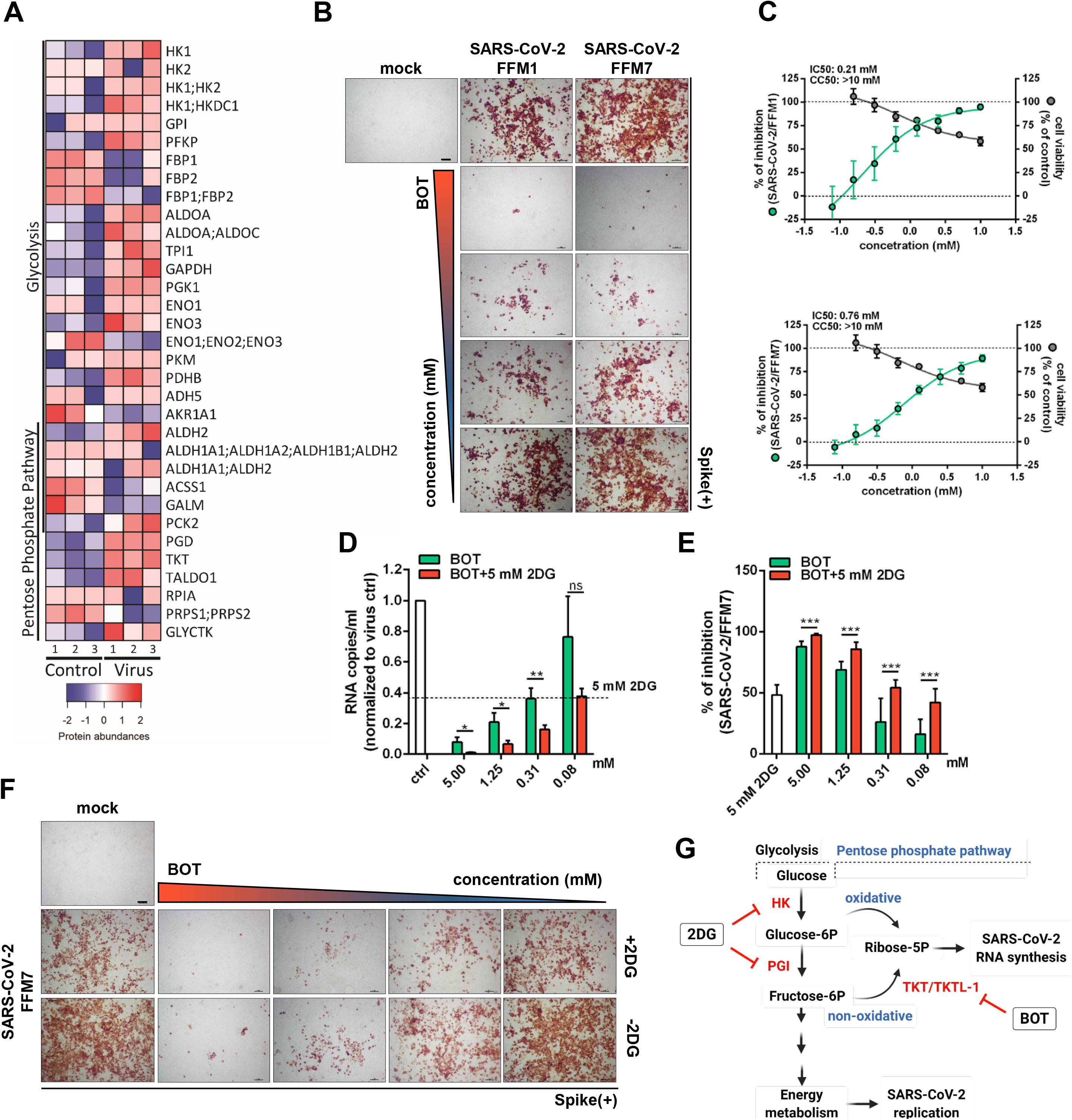
Glycolysis and pentose phosphate pathway as host target for antiviral therapy against SARS-CoV-2. **(A)** Heatmap of changes in protein abundance of components of glycolysis and pentose phosphate pathway in SARS-CoV-2 infected Caco-2 cells at 24h post infection. A Z score transformation was performed such that red and blue represent high and low protein abundance respectively. The plot was performed using the heatmaps2 function of the gplots package of the R suite. **(B)** Immunohistochemistry staining of SARS-CoV-2 spike protein in SARS-CoV-2/FFM1 and SARS-CoV-2/FFM7 infected Caco-2 cells treated with BOT. Caco-2 cells were pre-treated with different concentration of BOT for 24h. The cells were then infected with two different SARS-CoV-2 strains at MOI 0.01. 24h post infection, cells were fixed and stain for spike protein. Representative images of three independent experiments are shown. **(C)** Dose-response curves of viral inhibition and cell viability in BOT treated cells. Percentage of viral inhibition was evaluated by spike protein staining and cell viability was measured by MTT assay. The IC50 and CC50 values were determined using the curve regression function of GraphPad Prism 8. Both plots represent mean+SD of three independent experiments performed with three technical replicates. **(D)** Quantification of viral genomes in supernatant of SARS-CoV-2 infected Caco-2 cells treated with BOT in combination with 2DG or BOT alone. The number of SARS-CoV-2/FFM7 RNA was determined by qRT-PCR of RdRp gene and depicted as RNA copies/ml. The bar plot represents mean+SD of three independent experiments performed with three technical replicates. Statistical significance was determined with a two sided unpaired t-test. ns: not significant; * p≤0.05; ** p≤0.01. **(E)** Inhibition of viral infection in BOT treated cells in combination with 2DG. Caco-2 cells were pre-treated with different concentration of BOT for 24h. Then the 2DG at concentration 5mM was added and cells were infected with SARS-CoV-2/FFM7 at MOI 0.01. 24h post infection, cells were fixed and stain for spike protein. Percentage of viral inhibition was evaluated by spike protein staining. Bar graph depicts mean+SD of three independent experiments with three technical replicates. Statistical significance was determined with a two sided unpaired t-test. *** p≤0.005 **(F)** Immunohistochemistry staining of SARS-CoV-2 spike protein in SARS-CoV-2/FFM7 infected Caco-2 cells treated with BOT in combination with 2DG. Representative images of three independent experiments are shown. **(G)** Simplified scheme of glycolysis and pentose phosphate pathway. The targets for 2DG and BOT are depicted in red. The scheme was created with BioRender.com.

## Methods

### Proteomics data

The LC–MS/MS proteomics data with the dataset identifier PXD017710 have been previously published (Bojkova et al., 2020). Proteins derived from Caco-2 cells infected with SARS-CoV-2 after 24 h post infection and matching search terms “glycolysis” and “pentose phosphate pathway” in KEGG database were selected for further analysis. Significantly deregulated proteins (p value ≤ 0.5) in mock versus infected cells were plotted using the heatmaps2 function of the gplots package of the R suite. Briefly, raw enrichment values were used to calculate per row Z scores. Maximum and minimum Z score were attributed the colour red and blue respectively. Scripts available upon request.

### Cell culture and virus production

Caco-2 cells, a colon carcinoma derived cell line, was maintained in MEM (Minimal Essential Medium) containing 10% (v/v) foetal bovine serum,10,000 U penicillin/streptomycin and 2% (v/v) L-glutamine (Sigma Aldrich, Germany). Both SARS-CoV-2/FFM1 and SARS-CoV-2/FFM7 were isolated as previously described (Hoehl et al., 2020; Toptan et al., 2020). The viral stocks were produced by passaging virus on Caco-2 cells at MOI 0.1. Both strains underwent two passages. The virus titer was determined as TCID50.

### Antiviral and cytotoxicity assay

Caco-2 cells were seeded in 96-well plate. After reaching confluency, cell were pre-treated with benfooxythiamine (BOT, Zyagnum AG) for 24 h. Then the cells were additionally treated with 2-deoxy-D-glucose (2DG) or let untreated and infected with SARS-CoV-2 at MOI 0.01. 24 h later, cells and supernatants were harvested to determine viral load. The cytotoxic effect of BOT alone or in combination with 2DG was evaluated my MTT assay as previously described.

### Immunohistochemistry staining

Viral infection was assessed by staining of SARS-CoV-2 spike protein. 24h post infection, cells were fixed with acetone:methanol (40:60) solution followed by incubation with a primary monoclonal antibody directed against the spike protein of SARS-CoV-2 (1:1500, Sinobiological). Primary antibody was detected with a peroxidase-conjugated anti-rabbit secondary antibody (1:1000, Dianova), followed by addition of AEC substrate. The quantification of spike positive area was performed by BIOREADER®-7000-F-z-I (Bio-Sys).

### qRT-PCR of viral genome in supernatants

RNA from cell culture supernatant was isolated using the QIAamp Viral RNA Kit (Qiagen) according to the manufacturer’s instructions. Amount of viral RNA was detected by primers targeting the RNA-dependent RNA polymerase (RdRp): RdRP_SARSr-F2 (GTGARATGGTCATGTGTGGCGG) and dRP_SARSr-R1(CARATGTTAAASACACTATTAGCATA) using the Luna Universal One-Step RT-qPCR Kit (New England Biolabs) and a CFX96 Real-Time System, C1000 Touch Thermal Cycler. Number of viral copies was determined from standard curves generated by plasmid DNA (pEX-A128-RdRP) containing the corresponding amplicon regions of the RdRP target sequence.

### Statistics

All experiments were performed under similar conditions in three independent replicates. GraphPad Prism 8 software was used to plot the graphical representation and to perform statistical analysis. Statistical significance was calculated by two sided unpaired t-test as described in the figure legends (ns – not significant, * p < 0.05, ** p < 0.01, *** p < 0.005).

## References

Ayres, J.S. (2020). A metabolic handbook for the COVID-19 pandemic. Nat. Metab. 2, 572–585.

Bojkova, D., Klann, K., Koch, B., Widera, M., Krause, D., Ciesek, S., Cinatl, J., and Münch, C. (2020). Proteomics of SARS-CoV-2-infected host cells reveals therapy targets. Nature 583.

Codo, A.C., Davanzo, G.G., Monteiro, L. de B., de Souza, G.F., Muraro, S.P., Virgilio-da-Silva, J.V., Prodonoff, J.S., Carregari, V.C., de Biagi Junior, C.A.O., Crunfli, F., et al. (2020). Elevated Glucose Levels Favor SARS-CoV-2 Infection and Monocyte Response through a HIF-1α/Glycolysis-Dependent Axis. Cell Metab. 1–10.

Coy, J.F. (2017). EDIM-TKTL1/Apo10 blood test: An innate immune system based liquid biopsy for the early detection, characterization and targeted treatment of cancer. Int. J. Mol. Sci. 18, 1–18.

Deng, Y., Lei, L., Chen, Y., and Zhang, W. (2020). The potential added value of FDG PET/CT for COVID-19 pneumonia. Eur. J. Nucl. Med. Mol. Imaging 47, 1634–1635.

Shen, B., Yi, X., Sun, Y., Bi, X., Du, J., Zhang, C., Quan, S., Zhang, F., Sun, R., Qian, L., et al. (2020). Proteomic and Metabolomic Characterization of COVID-19 Patient Sera. Cell 59–72.

Stincone, A., Prigione, A., Cramer, T., Wamelink, M.M.C., Campbell, K., Cheung, E., Olin-Sandoval, V., Grüning, N., Krüger, A., Tauqeer Alam, M., et al. (2015). The return of metabolism: biochemistry and physiology of the pentose phosphate pathway. Biol. Rev. 90, 927–963.

Thomas, T., Stefanoni, D., Reisz, J.A., Nemkov, T., Bertolone, L., Francis, R.O., Hudson, K.E., Zimring, J.C., Hansen, K.C., Hod, E.A., et al. (2020). COVID-19 infection alters kynurenine and fatty acid metabolism, correlating with IL-6 levels and renal status. JCI Insight 5.

Tylicki, A., Lotowski, Z., Siemieniuk, M., and Ratkiewicz, A. (2018). Thiamine and selected thiamine antivitamins — biological activity and methods of synthesis. Biosci. Rep. 38, 1–23.

Zhao, J., and Zhong, C.J. (2009). A review on research progress of transketolase. Neurosci. Bull. 25, 94–99.

Zhu, L., She, Z.-G., Cheng, X., Qin, J.-J., Zhang, X.-J., Cai, J., Lei, F., Wang, H., Xie, J., Wang, W., et al. (2020). Association of Blood Glucose Control and Outcomes in Patients with COVID-19 and Pre-existing Type 2 Diabetes. Cell Metab. 31, 1068–1077.e3.

## References

Hoehl, S., Rabenau, H., Berger, A., Kortenbusch, M., Cinatl, J., Bojkova, D., Behrens, P., Böddinghaus, B., Götsch, U., Naujoks, F., et al. (2020). Evidence of SARS-CoV-2 infection in returning travelers from Wuhan, China. N. Engl. J. Med. 382, 1278–1280.

Toptan, T., Hoehl, S., Westhaus, S., Bojkova, D., Berger, A., Rotter, B., Hoffmeier, K., Cinatl, J., Ciesek, S., and Widera, M. (2020). Optimized qRT-PCR Approach for the Detection of Intra- and Extra-Cellular SARS-CoV-2 RNAs. Int. J. Mol. Sci. 21, 4396.

